# Direct evidence from kinetic D_2_O-MRI modeling that the choroid plexus is not the major source of cerebrospinal fluid

**DOI:** 10.64898/2026.01.20.700750

**Authors:** Zhao Zhang, Yun Wu, Yanmin Zheng, Ye Li, Qiong Ye

## Abstract

The role of the choroid plexus (CP) as the primary source of cerebrospinal fluid (CSF) remains controversial. Here, we used dynamic indirect D_2_O magnetic resonance imaging (MRI) at 9.4T to investigate whole-brain CSF dynamics in rats. Spin-echo echo-planar imaging (SE-EPI) was performed during intravenous D_2_O infusion with a temporal resolution of 10.69 s and an in-plane resolution of 150 × 150 µm. An echo time of 150.0 ms was employed to effectively suppress signals from brain parenchyma, blood, and interstitial fluid, resulting in preferential visualization of CSF. The ambient and supracerebellar cisterns (AC+SC) exhibited the greatest signal attenuation, reaching 47.32 ± 15.70% of baseline, whereas the lateral ventricles (LV1, LV2) showed smaller reductions (68.73 ± 18.72%–72.64 ± 15.07% of baseline). CSF production rates were quantified using a one-compartment perfusion model. Estimated secretion rates were 0.11 and 0.13 µL/min in the lateral ventricles and 1.30 µL/min in the AC+SC region, indicating a 5.42-fold higher CSF production outside the ventricles. These findings provide direct in vivo evidence that the lateral ventricles are not the primary site of CSF production. The proposed D_2_O-based dynamic SE-EPI approach is safe, non-invasive, cost-effective, and suitable for widespread application.

## Introduction

The secretion of cerebrospinal fluid (CSF) has become a significant area of investigation in recent years, with a growing recognition of the factors that influence CSF movement in the central nervous system (CNS), e.g., sleep, age, Alzheimer’s disease (AD), cardiac rhythms[1-7]. The secretion of CSF is a key mechanism in maintaining the fluid balance within the CNS. As proposed by the “third circulation” theory, CSF is produced by the choroid plexus (CP) located in the roofs of the lateral, third, and fourth ventricles. Once produced, CSF flows through the ventricular foramina and fills the subarachnoid space (SAS), surrounding both the brain and spinal cord. Yet, the exact sources of CSF production remain controversial. In an early experiment, the complete surgical removal of the CP did not eliminate CSF production[8]. Extrachoroidal sources, including the filtration or secretion of fluid across capillary walls, may also contribute to the overall production of CSF [9-11]. A study reported no significant differences in signal changes in CSF between AQP-1 knockout and wild-type mice, using intravenous administrated H_2_^17^O, while AQP-4 knockout mice demonstrated a significant reduction in water influx into the CSF space[12]. The CSF AQP-1 and AQP-4 are predominantly expressed in the choroid plexus epithelium and in the subpial and perivascular endfeet of astrocytes, respectively[12].

Moreover, the secretion of CSF can be influenced by multiple factors, such as age, sex, anesthesia protocol, AD, and vasopressin levels [13-15]. Previous studies have employed a wide range of direct or indirect methods to quantify the rate of CSF production[16, 17], but the results vary significantly. The review by Guojun Liu et al.[18] provided a thorough summary of this topic. To date, no method has been established that is safe, accurate, and reproducible.

MRI is a non-invasive technique that provides detailed structural, functional, and physiological information, making it ideal for studying CSF production. Previous studies have used phase-contrast magnetic resonance imaging (PC-MRI)[19-21] and arterial spin labeling (ASL-MRI) for non-invasive detection of CSF production[13-15], but these methods have limitations, such as stringent assumptions, low spatial resolution, and the fact that they only target CSF production from the lateral ventricles.

An alternative is isotopic water MRI, including D_2_O and H_2_^17^O[16, 22-24]. Both are commercially available, stable and non-radioactive. Direct imaging is challenged by low sensitivity due to the lower gyromagnetic ratios of D and O-17 (6.54 MHz/T and 5.77 MHz/T, respectively)[16]. Moreover, specific RF coils are required, which makes indirect imaging preferable. In this approach, the principle of signal attenuation by H_2_^17^O relies on the reduction of T_2_ or T_1_ρ relaxation of protons in bulk water [23, 25, 26], while the attenuation of the signal by D_2_O primarily results from the decrease in proton density due to displacement [16, 25].

As D_2_O is administered intravenously, the molecules perfuse into the tissue and partially replace H_2_O in the tissue, thereby reducing proton density and attenuating the signal intensity in ^1^H MRI due to the replacement effect of isotopic substitution[16]. The intravenous use of D_2_O is safe, except at very high doses over the long term[27]. Previous studies have demonstrated the safety of doses in rats at 2 ml/100 g[16] and 1.5ml/100 g[28]. An initial clinical trial with a dose of 0.43 ml/kg proved its safety in human[29]. Furthermore, the chemical and physical properties of D_2_O are similar to those of regular water molecules, allowing it to pass through aquaporin[27]. Compared to H_2_^17^O, D_2_O is more affordable (around $100 per 100 ml), and its quantification is more straightforward. Based on all of the above-mentioned advantages, indirect D_2_O MRI is an ideal candidate for the study of CSF production.

Indirect D_2_O MRI with intravenous administration of D_2_O, aimed at understanding CSF production, has been reported very recently [28]. However, the temporal resolution of 4:21 minutes per phase with two phases, pre- and post-infusion, is insufficient to distinguish between choroidal and extrachoroidal sources of CSF production. Additionally, several previous studies used long echo time (TE) to eliminate signals from brain parenchyma, blood, and interstitial fluid (ISF), obtaining excellent, exclusive visualization of CSF due to its significantly longer transverse relaxation time[30-32]. Our study used high spatiotemporal resolution imaging with long TE during continuous intravenous contrast agent infusion to provide direct evidence for the sources of CSF production. Kinetic modeling was performed to extract quantitative information on the rates of CSF production in choroidal and extrachoroidal sources. Our approach has several advantages: it is non-invasive, safe, requires no additional hardware or software, is cost-effective, straightforward in post-processing, and can yield quantitative results.

## Materials and methods

All animal experiments were approved by the local institutional animal experiment committee.

### MRI scan

The MRI data were acquired using a 9.4T scanner (Bruker, Ettingen, Germany) equipped with a home-made transceiver rat brain surface coil. Eleven normal adult female Sprague-Dawley rats (195.3-322.0 g, 2-4 months of age) were used in this study. Animals were anesthetized with 3-4% isoflurane for induction and maintained with 1-2.5% isoflurane during the scan. A normal saline-filled tubular phantom was positioned on top of the brain as a reference. The tail vein was cannulated with a 26G IV catheter and connected to an infusion pump via PE-50 tube. A water heating pad was used to maintain body temperature. Respiratory rate was continuously monitored and maintained at 30–60 breaths per minute.

As illustrated in Fig 1, after a localizer scan for positioning, the scan protocol sequentially included these sequences.

**Figure 1.**
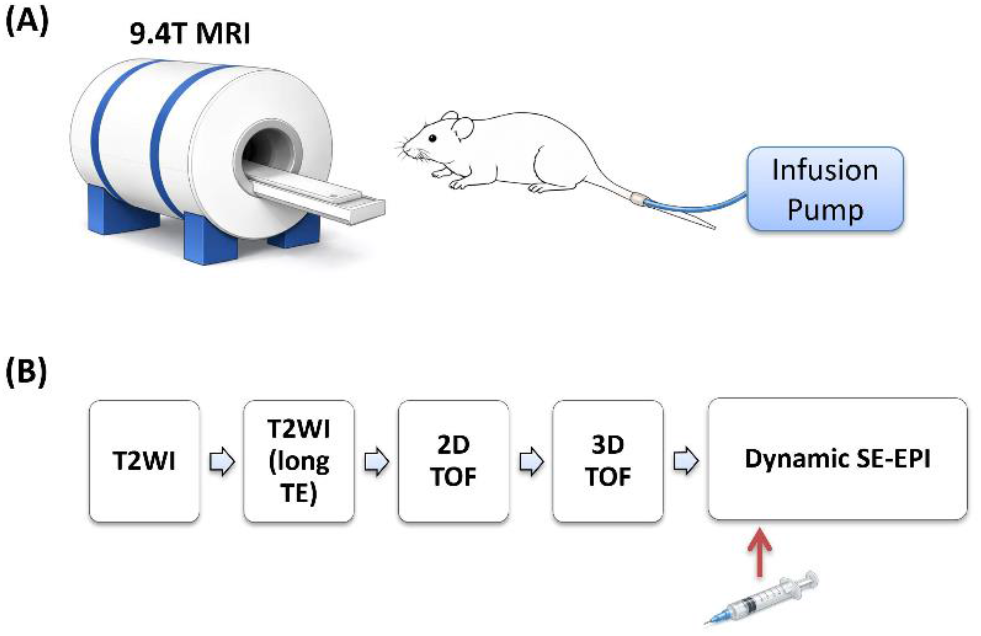
Schematic illustration of D_2_O MRI for the investigation of cerebrospinal fluid (CSF) production. (A) Rats were cannulated via the tail vein using an intravenous catheter, and D_2_O was administered through an infusion pump. MRI acquisition was performed at 9.4 T during continuous D_2_O infusion. (B) Detailed MRI sequences included in the experimental protocol. D_2_O infusion was initiated at the end of the fifth phase of dynamic spin-echo echo planer imaging (SE-EPI).

T2WI image was acquired for the anatomy with these parameters: TE/Repetition time (TR)=24.0/5000.0 ms, Echo spacing=8.0 ms, Rare factor=8, 2D axial slices, FOV=27×27 mm^2^, matrix=256×256, slice thickness=0.5mm without gap, No. of slices=60, resolution=0.105×0.105 mm^2^, Number of averages (NA)=4, scan time=10min40sec.

T2WI image with long TE was scanned for the visualization of CSF: TE/TR=120.0/11160.0 ms, Echo spacing= 40.0 ms, RARE factor=8, 2D axial slices, FOV=20×16 mm^2^, matrix=200×160, slice thickness=1.0mm, without gap, No. of slices=30, resolution=0.1×0.1 mm^2^, NA=4, scan time=14min52sec800ms.

2D time-of-flight (TOF) for the anatomy of cerebral vessels: TE/TR=1.9/12.0 ms, flip angle=80°, 2D axial slices, FOV=27×27 mm^2^, matrix=256×256, slice thickness=0.5mm, No. of slices=60, resolution=0.105×0.105 mm^2^, NA=2, scan time=6min8sec640ms.

3D TOF for the anatomy of cerebral vessels: TE/TR=1.8/12.0 ms, flip angle=20, 3D axial slices, FOV=27×27×30 mm^3^, matrix=256×256×96, resolution=0.105×0.105×0.313 mm^2^, NA=2, scan time=8min8sec448ms.

Dynamic single-shot spin-echo echo planer imaging (SE-EPI) was acquire to capture the dynamics of CSF production: TE/TR=150.0/3562.0 ms, Δt=10sec686ms (temporal resolution), Spin Echo, FOV=22×16 mm^2^, matrix=147×107, slice thickness= 1.5mm, No. of slices=20, voxel=0.15×0.15 mm^2^, 2D axial slices, NA=3, repetitions=120, scan time=21min22sec320ms. A dose of 2 mL/100 g of 0.9% saline D_2_O was administered using an infusion pump at a rate of 1 mL/min. The injection was initiated after five repetitions to ensure a stable baseline.

Both 2D TOF and 3D TOF images were acquired because of their different sensitivities to arteries and veins.

### Post-processing and image analysis

All images were converted to NIfTI format using Bruker2NIfTI. Brain extraction was performed in 3D Slicer (version 5.10).

Four-dimensional dynamic SE-EPI images were motion-corrected in Fiji (NIH, USA). Averaged images were generated for region-of-interest (ROI) delineation. Two-dimensional ROIs were manually placed on the lateral ventricles (LV1 and LV2), third ventricle (3V), fourth ventricle (4V), cerebral aqueduct (Aq), ambient and supracerebellar cisterns (AC+SC), and dorsal frontal cistern (DFC). Dynamic signal changes were extracted from each ROI. ROI volumes were calculated by multiplying the number of included pixels by the volume of pixel.

The 3D volumes of the CSF compartments were derived from T2WI with long TE. Due to the relatively low spatial resolution (0.1 × 0.1 × 1.0 mm^3^), only the 3D volumes of LV1, LV2, and AC+SC were calculated. The binary CSF masks were generated using the Triangle method as a reference. The 3D ROIs for LV1, LV2, and AC+SC were manually delineated on the corresponding consecutive slices. The 3D volumes of LV1, LV2, and AC+SC were calculated by multiplying the number of pixels by the volume of pixel.

### Kinetic modeling of dynamic SE-EPI D_2_O imaging

The first point of dynamic images was excluded from further analysis. The 2^nd^ to 5^th^ dynamic images were acquired before injection, and were used as baseline signal. Normalized signal relative to baseline (%), *r*(*t*), were calculated.

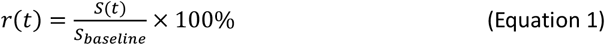

*S*(*t*), the dynamic signal intensity. *S*_*baseline*_, the signal intensity at baseline.

Signal Change (%) relative to baseline, *S*^′^(*t*), were calculated.

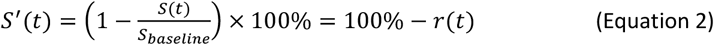

To quantify the production rate of CSF, a single-compartment perfusion model was applied[33], assuming equal production and clearance rates.

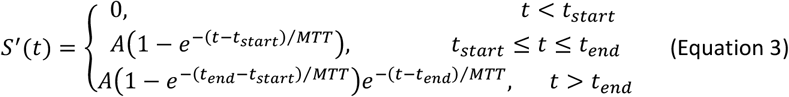

A, Plateau-phase maximum signal change (%). MTT: Mean transit time in the ROI (min). The *t*_*start*_ and *t*_*end*_ are the time points at the start of signal change and the peak of signal intensity. The production rate of CSF, *Q*_*CSF*_, can be calculated by

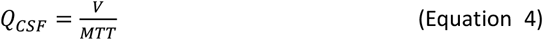

V, the volume to 2D ROI. Lsqcurvefit was used for curve fitting.

The derived *Q*_*CSF,corrected*_ was corrected by the volumes of 2D ROI and 3D ROI.

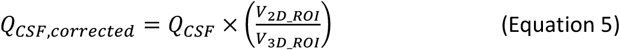

### Statistics

Statistical analyses were performed in MATLAB (R2022b). Mean ± standard deviation (STD) were calculated for the 3D volumes of LV1, LV2, and AC+SC. Group-averaged dynamic SE-EPI signal intensities and corresponding STDs were obtained for all analyzed CSF compartments.

## Results

### The anatomical relationships among brain parenchyma, CSF, and blood vessels

The relative positions of the brain parenchyma, CSF, and blood vessels were visualized using ITK-SNAP (NIH, USA) based on fused images of T2WI, T2WI with long TE, and 2D TOF imaging (Figure 2). As indicated by the yellow arrows, the subarachnoid space (SAS) adjacent to the superior sagittal sinus can be clearly visualized [34]. As shown in Figure 3, three-dimensional reconstructions derived from long-TE T2WI and 3D TOF imaging provide a comprehensive view, allowing improved visualization of the anatomical relationships among these structures.

**Figure 2.**
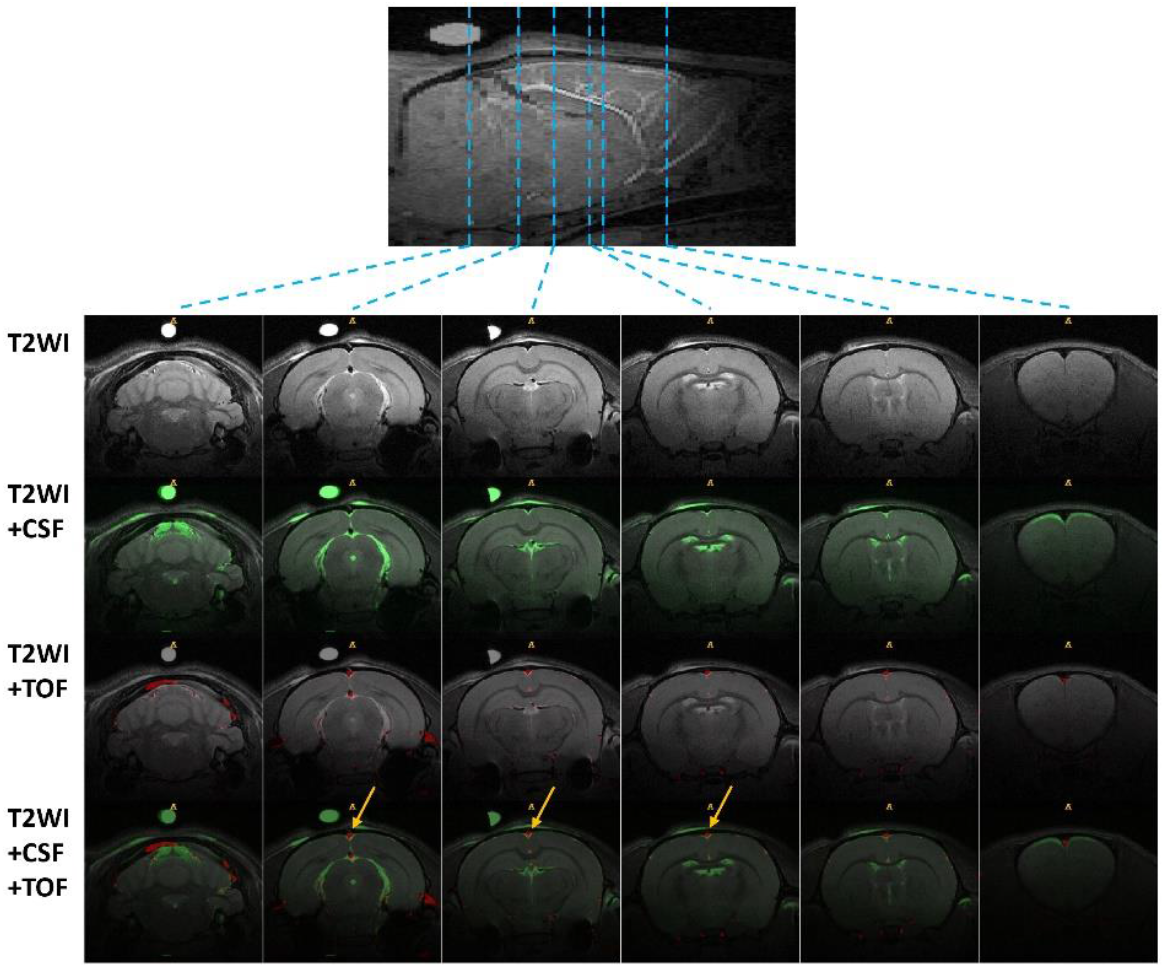
Sagittal anatomical image with slice positions indicated. T2WI shows the anatomy of the corresponding slices. T2WI+CSF represents the fused images of T2WI and T2WI with long TE, highlighting the distribution of CSF. T2WI+TOF refers to the fused images of T2WI and 2D TOF, emphasizing the vasculature. T2WI+CSF+TOF combines T2WI, T2WI with long TE, and 2D TOF, illustrating the anatomical relationship between brain parenchyma, CSF, and blood vessels. As indicated by the yellow arrows, the subarachnoid space adjacent to the superior sagittal sinus is clearly visible. T2WI, T2-weighted imaging; CSF, cerebrospinal fluid; TE, echo time; TOF, time-of-flight.

**Figure 3.**
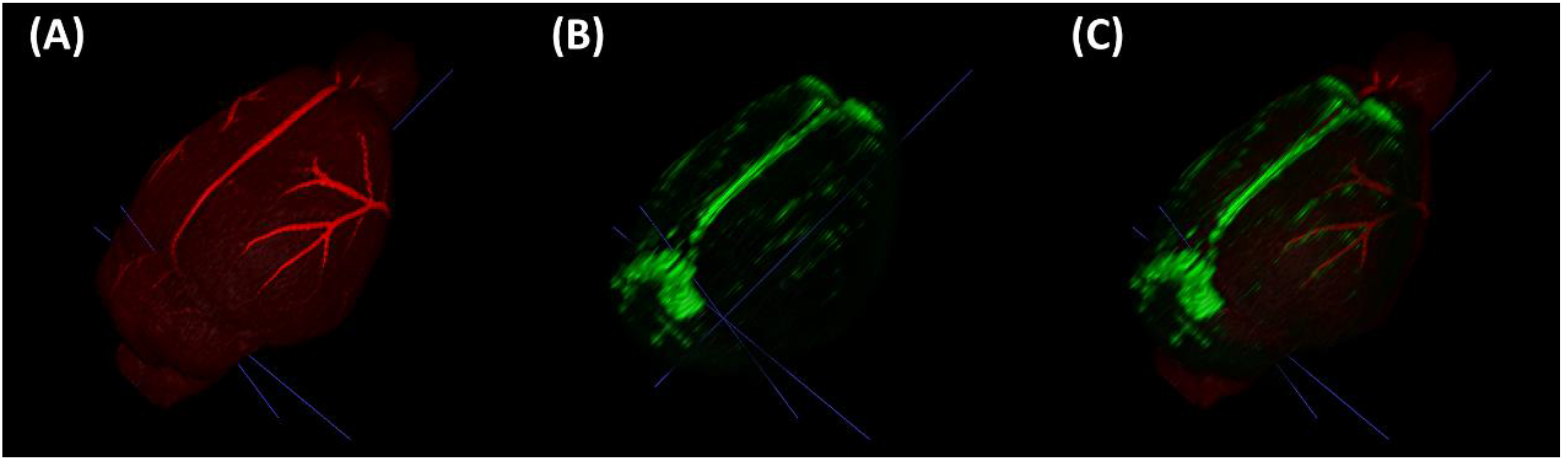
(A) 3D rendering of 3D TOF imaging showing the blood vessels on the surface of the brain. (B) 3D rendering of CSF on the brain surface imaged by T2WI with long TE. (C) Merged view of (A) and (B), providing a clear visualization of the anatomical relationship between the blood vessels and CSF on the brain surface. TOF, time-of-flight; CSF, cerebrospinal fluid; T2WI, T2-weighted imaging; TE, echo time.

### Dynamic D_2_O SE-EPI images and quantitative dynamic curves

As shown in the upper row of Figure 4, signal intensities in the lateral ventricles (LV1 and LV2) exhibited a moderate decline following D_2_O administration, whereas signal intensity in the third ventricle and the suprasellar cistern remained relatively stable. In the lower row of Figure 4, at the level of the ambient cistern, signal intensity in the AC+SC decreased markedly, while only subtle changes were observed in the cerebral Aq.

**Figure 4.**
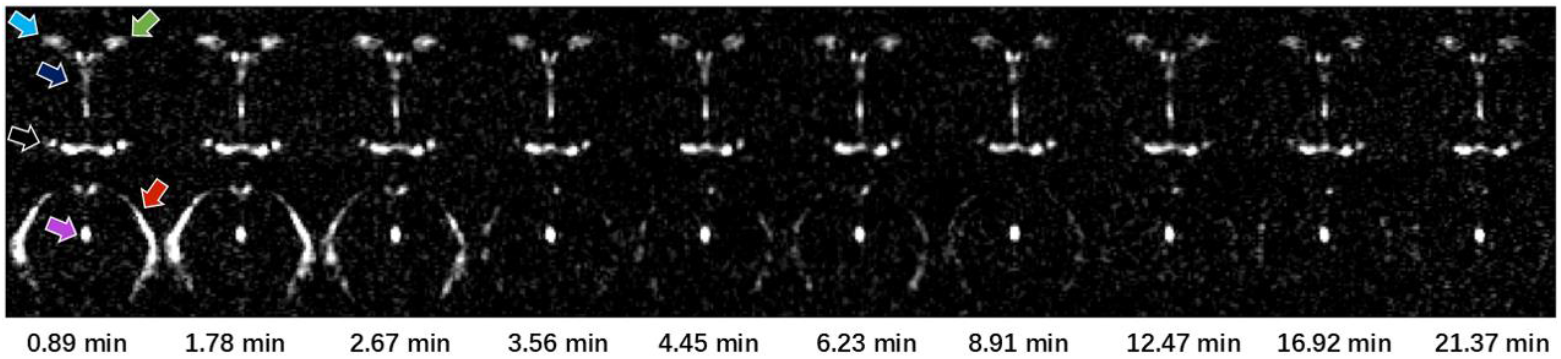
Dynamic D_2_O-induced signal attenuation in a representative rat brain (not continuous phases). The upper row shows the slice at the level of the lateral ventricles, where light green and light blue arrows indicate the lateral ventricles (LV1, LV2), the dark blue arrow indicates the third ventricle (3V), and the black arrow points to the suprasellar cistern. The lower row shows the slice at the level of the ambient and supracerebellar cisterns (AC+SC), with the red arrow indicating the AC+SC, and purple arrow pointing to the cerebral aqueduct (Aq). The first column represents the final phase before the infusion of D_2_O. Signal intensities in the AC+SC, LV1, and LV2 exhibit a notable decrease over time, while other regions show minimal change.

Phases 2–5 acquired prior to D_2_O infusion were defined as the baseline. Signal intensities were normalized to this baseline for each rat, and the group-averaged normalized signal intensity (%) is presented in Figure 5. A delay of approximately 0.71 min (∼43 s) was observed between D_2_O infusion and the onset of signal change.

**Figure 5.**
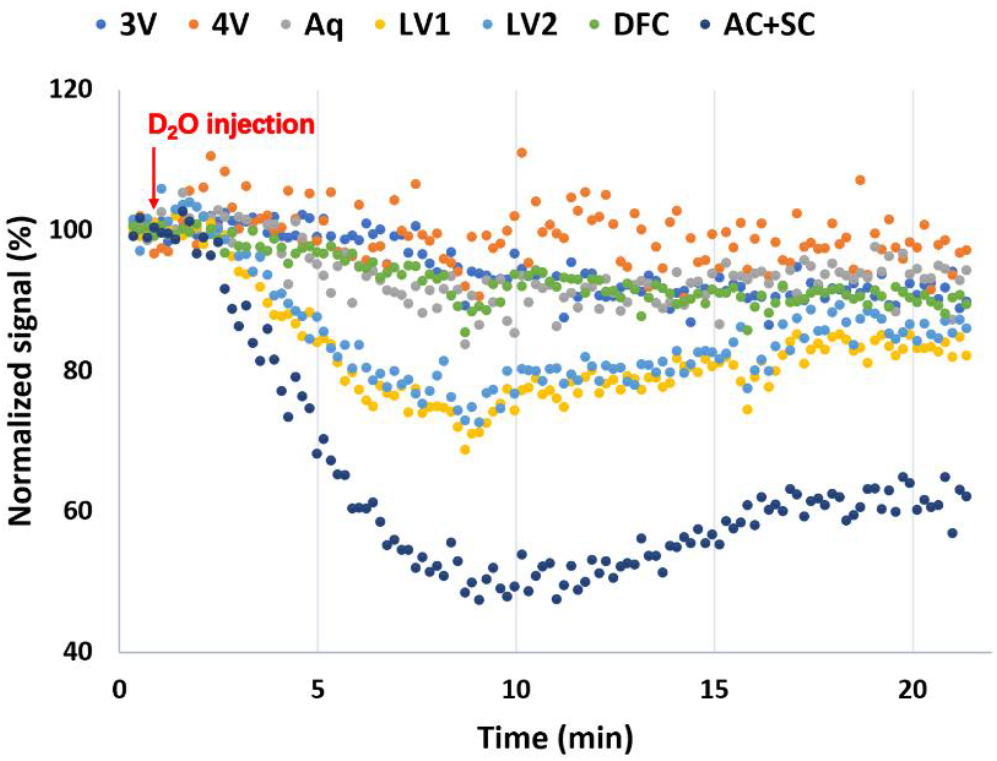
Group-averaged normalized signal (% of baseline) time course in the CSF compartments.

Among all analyzed regions, AC+SC exhibited the most pronounced signal reduction, reaching a minimum of 47.32 ± 15.70% of baseline at 9.08 min, followed by a moderate recovery. LV1 and LV2 showed nearly identical dynamic profiles, with signal intensities decreasing to 68.73 ± 18.72% at 8.73 min and 72.64 ± 15.07% at 9.08 min, respectively, followed by partial recovery. The regional dynamic curves with mean ± SD are provided in the supplementary materials (S1). In contrast, the 3V, 4V, Aq, and DFC showed a gradual and sustained signal decline without evident recovery.

### Kinetic modeling of dynamic D_2_O SE-EPI images

A single-compartment perfusion model was employed to quantify the CSF production rate. The results of the curve fitting are presented in Fig. 6, with the fitted MTT and *Q*_*CSF*_ values listed in Table 1. Since only LV1, LV2, and AC+SC had measured 3D volumes (2.12 ± 0.68 µL, 2.33 ± 0.87 µL, and 38.37 ± 10.34 µL, respectively), the *Q*_*CSF*_ was corrected for these regions. The *Q*_*CSF,corrected*_ values were 0.11, 0.13, and 1.30 µL/min for LV1, LV2, and AC+SC, respectively, with corresponding MTT values of 18.87, 17.66, and 29.48 minutes. The CSF production rate in the AC+SC region was approximately 5.42 times higher than that in the lateral ventricles (LV1+LV2).

**Table 1.** The derived results of the CSF production rate.

| *Name of ROI* | *volume of ROI (uL)* | *MTT (min)* | *$Q_{CSF}$ (uL/min)* | *3D volume (uL)* | *$Q_{CSF,corrected}$ (uL/min)* |
| --- | --- | --- | --- | --- | --- |
| **3V** | 2.64 | 1.20E+05 | 2.20E-05 |  |  |
| **4V** | 1.01 | 4.53E+04 | 2.22E-05 |  |  |
| **Aq** | 2.16 | 23.82 | 0.09 |  |  |
| **LV1** | 1.96 | 18.87 | 0.10 | 2.12 | 0.11 |
| **LV2** | 1.77 | 17.66 | 0.10 | 2.33 | 0.13 |
| **DFC** | 8.66 | 1.07E+05 | 8.09E-05 |  |  |
| **AC+SC** | 9.35 | 29.48 | 0.32 | 38.37 | 1.30 |

**Figure 6.**
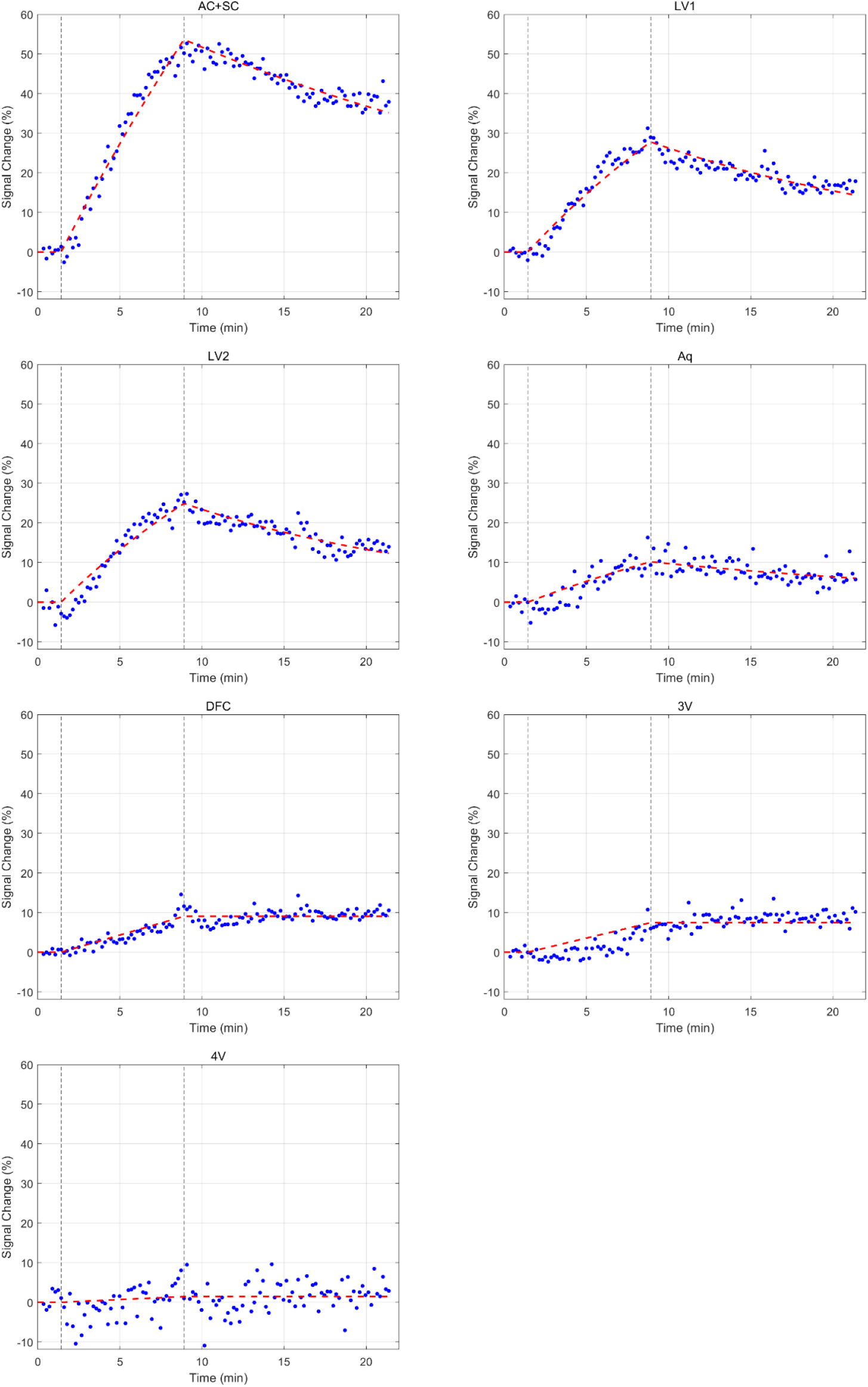
Kinetic modeling for quantifying CSF production.

## Discussion

Our study provides direct quantitative evidence that the choroid plexus is not the primary site of CSF production. We developed a noninvasive in vivo approach that combines a single intravenous dose of D_2_O with high spatiotemporal-resolution SE-EPI MRI at long TE. This method ensures a robust SNR and superior CSF visualization. Notably, T2WI with long TE also can visualize the SAS adjacent to the superior sagittal sinus. When combined with vascular imaging, this technique allows for an integrated examination of both the CSF system’s structure and function, as well as its spatial relationship with the cerebral vasculature. This study offers a novel approach for investigating factors influencing CSF production and holds significant potential for advancing research on the glymphatic system. Furthermore, this protocol can be performed on all conventional MRI scanners without the need for additional hardware or specialized sequences, and the quantification process is relatively straightforward. This technology can be widely implemented.

The D_2_O MRI has been developed for decades[16, 22]. Direct imaging requires specialized deuterium coils and suffers from a lower SNR, whereas indirect proton-based imaging detects the reduction in proton density caused by D_2_O replacement, thus eliminating the need for dedicated hardware and sequences[16, 25, 35]. Therefore, this study employed an indirect imaging approach. A 2D multi-slice SE-EPI sequence was used, offering rapid acquisition, high SNR, and relative insensitivity to magnetic field inhomogeneity and motion. The protocol achieved a temporal resolution of 10.69 sec, a slice thickness of 1.5 mm, and an in-plane spatial resolution of 150 × 150 µm, enabling high spatiotemporal dynamic imaging. Furthermore, the use of a long TE effectively suppressed signals from blood, brain parenchyma, and interstitial fluid, allowing selective visualization of CSF. This methodology may provide the most direct evidence to date regarding the mechanisms of CSF production.

In dynamic SE-EPI imaging, newly secreted CSF in the lateral ventricles exhibited an initial increase followed by decay, whereas the changes in 3V, 4V, and Aq were markedly slower, indicating substantial diversion of CSF from the classical ventricular pathway. Potential routes include perivascular spaces between the lateral ventricles and the subarachnoid space [36] or direct absorption into the brain parenchyma[37]. The finding aligns with the work by Marin Bulat et al.[38], where 3H-water, slowly injected into the lateral ventricle, was rapidly absorbed by the periventricular capillaries rather than flowing into the subarachnoid space. Consistent with this, the DFC, which connects to the SAS, shows signal dynamics nearly identical to those of 3V, 4V, and Aq, may further supporting a lateral ventricular origin for CSF in this region.

Notably, the CSF production rate in the AC+SC region was approximately five times higher than that in the lateral ventricles, although its precise source remains unclear. Considering the spatial relationship between CSF compartments and the intracranial vasculature, we hypothesize that the pial arterial network at the cerebrum–cerebellum interface, rather than large venous structures such as the superior sagittal sinus, is the more likely site of CSF generation. This interpretation is further supported by the slow signal kinetics observed in the DFC, which lies along the same venous drainage route but shows minimal dynamic change, suggesting that venous structures contribute minimally to CSF production. Additionally, Sweet and Locksley provided evidence of bidirectional exchange of water and electrolytes between blood and cerebrospinal fluid across both the ventricular and subarachnoid compartments [39].

In rats, total ventricular CSF production was measured at 3.38 μL/min using an indirect tracer dilution method with CSF sampled from the cisterna magna[40], whereas direct sampling from both lateral ventricles and the third ventricle yielded production rates ranging from 0.39 to 1.40 μL/min [41]. Our result, 1.54 μL/min derived from the summed quantitative analysis of LV1, LV2, and AC+SC, aligns in magnitude with these previous findings. Yet CSF production rate is influenced by multiple factors. In accordance with prior studies, our research also observed considerable inter-individual variability (supplemental material, S1).

This study has several limitations, including the fact that the signal changes in CSF reflect both proton density replacement by D_2_O and relaxation effects. As reported by Fu-Nien Wang et al.[16], the replacement effect is the dominant factor in in vivo perfusion imaging, and the relaxation effects were not corrected in this study. Additionally, while the administered contrast agent dose is generally considered safe, such supraphysiological loading may transiently alter hemodynamics and electrolyte balance, factors that should be considered when interpreting the findings.

Future studies incorporating a dual-inversion pulse into the sequence could further suppress non-CSF signals, resulting in a purer CSF contrast with improved SNR efficiency. Additionally, combining multi-channel coils with compressed sense techniques is expected to significantly accelerate imaging. Achieving higher SNR efficiency and reducing partial volume effects in future studies will enable pixel-wise kinetic modeling at even higher spatial resolutions.

### Conclusion

Our D_2_O-based dynamic SE-EPI imaging during the intravenous infusion approach provides direct evidence that the lateral ventricles are not the primary site of CSF production, as the CSF secretion rate in the AC+SC region is substantially higher than that in the lateral ventricles.

## Acknowledgements

We thank the 9.4 T High Field Magnetic Resonance Imaging System in Superconducting Magnet SM4 of the Steady High Magnetic Field Facility, CAS for the assistance on the experiment (https://cstr.cn/31125.02.SHMFF.SM4.MRI). All authors would like to thank Hongyi Yang, Ganghan Yang, and Zhuoyang Lin for their support in coil testing, Dr. Qingwang Liu for providing the test rat, Dr. Wei Tong and Dr. Tingting Shao for their discussions on the experiments, and Mr. Qingping Wu from the Asset Department for his non-technical assistance.

## Data Availability Statement

Data and code will be made available upon reasonable request after submitting a formal project outline.

