## Supplemental figure S1 for "Direct evidence from kinetic D_2_O-MRI modeling that the choroid plexus is not the major source of cerebrospinal fluid"

**Supplemental material**

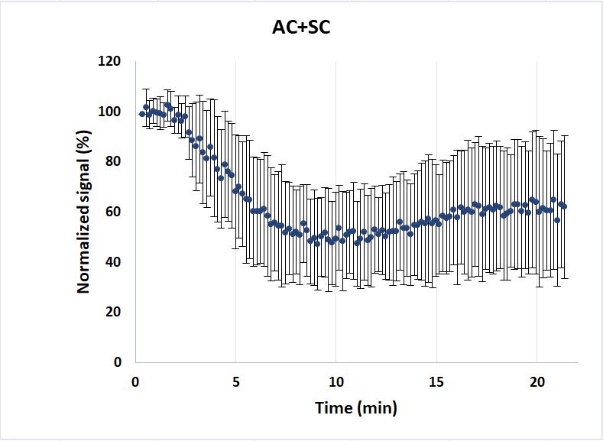

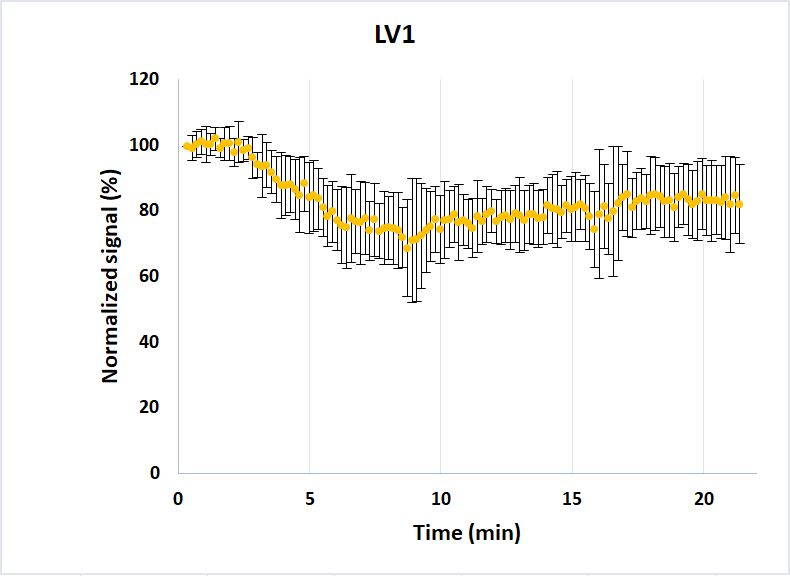

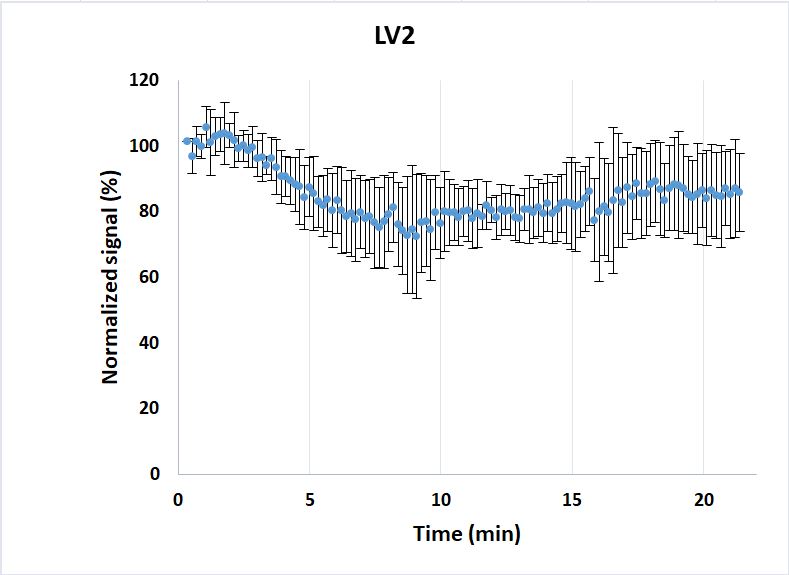

S1. **Group-averaged normalized signal (% of baseline) time course in the AC+SC, LV1, and LV2 (n=11).** Data are presented as mean ± standard deviation (STD). AC+SC, ambient and supracerebellar Cisterns; LV1, LV2, lateral ventricles.
